# Feature selection with vector-symbolic architectures: a case study on microbial profiles of shotgun metagenomic samples of colorectal cancer

**DOI:** 10.1101/2024.11.18.624180

**Authors:** Fabio Cumbo, Simone Truglia, Emanuel Weitschek, Daniel Blankenberg

## Abstract

The continuingly decreasing cost of next-generation sequencing has recently led to a significant increase in the number of microbiome-related studies, providing invaluable information for understanding host-microbiome interactions and their relation to diseases. A common approach in metagenomics consists of determining the composition of samples in terms of the amount and types of microbial species that populate them, with the goal to identify microbes whose profiles are able to differentiate samples under different conditions with advanced feature selection techniques. Here we propose a novel backward variable selection method based on the hyperdimensional computing paradigm, which takes inspiration from how the human brain works in the classification of concepts by encoding features into vectors in a high-dimensional space. We validated our method on public metagenomic samples collected from patients affected by colorectal cancer in a case/control scenario, by performing a comparative analysis with other state-of-the-art feature selection methods, obtaining promising results.

**AUTHOR SUMMARY:** Characterizing the microbial composition of metagenomic samples is crucial for identifying potential biomarkers that can distinguish between healthy and diseased states. However, the high dimensionality and complexity of metagenomic data present significant challenges in the context of accurately selecting features. Our backward variable selection method, based on the hyperdimensional computing paradigm, offers a promising approach to overcoming these challenges. By effectively reducing the feature space while preserving essential information, this method enhances the ability to detect critical microbial signatures associated with diseases like colorectal cancer, leading to more precise diagnostic tools.

## INTRODUCTION

Over the last decade, the cost of next-generation sequencing technologies has steadily decreased, a trend that is expected to continue. This reduction has positively impacted all areas of the life sciences, resulting in the generation of substantial biomedical data. Metagenomics, which involves the study of microorganisms in biological samples and their relationship to pathological conditions and environmental factors, is one field that has particularly benefited from these advancements. Metagenomics frameworks that rely on sophisticated statistical analysis and machine learning techniques allow the identification of relevant signatures across a wide range of pathologic conditions by inspecting differences in the abundance of specific microbial species in large cohorts of samples in a case/control scenario. Many studies have demonstrated the involvement of microbes in the genesis and development of a multitude of health-related conditions, including cancer.

Here we focus in particular on the disease of colorectal cancer (CRC), which is the second most common cause of cancer death worldwide (916,000 deaths) and the third for number of cases per year (1.93 million cases in 2020) according to the World Health Organization [1]. Additionally, the American Cancer Society (ACS) indicates that the 5-year survival rate (i.e., percentage of people affected by the same type and stage of cancer still alive after 5 years from the first diagnosis) is approximately 65% when the cancer is detected at an early stage [2]. Therefore, noninvasive methods allowing early detection of CRC are crucial to enhancing patient survival. Numerous studies have recently examined the difference in the abundance of specific microbes in metagenomics samples (usually fecal samples in the case of CRC) collected from case and healthy individuals in order to establish whether it can be used as a fast, accurate, and noninvasive tool for diagnosing the disease [3].

Here we propose a novel stepwise feature selection method following a backward variable elimination strategy based on the brain-inspired hyperdimensional computing paradigm. Hyperdimensional (HD) computing is a relatively new computational paradigm that aims to replicate the structure and function of the brain in order to perform various computational tasks. It is based on the idea that the brain uses distributed representations to encode and process information which is represented by unique vectors in a high-dimensional space [4–6]. These vectors are often referred to as “semantic pointers” because they are thought to encode the meaning of the information they represent [7].

This technology has been recently adopted to perform various computational tasks such as classification, clustering, and pattern recognition in different scientific domains [8–12]. One of the key benefits of HD computing is its efficiency because vectors used to represent information are distributed throughout the high-dimensional space, making it possible to perform many operations in parallel, which makes it well-suited for handling large amounts of data. It is brain-inspired in the sense that every concept is encoded into vectors in a high-dimensional space that can be combined together with simple arithmetic operations in order to represent more complex concepts, in the same way the human brain works with the associative memory for distinguishing very similar concepts by remembering patterns [4,13,14]. We integrated our method as a subroutine of *chopin2* [15,16], an open-source Python 3.8 package originally developed for classifying massive datasets with commodity hardware.

To the best of our knowledge, this is the first attempt of building a feature selection method following this emerging computing paradigm.

## RESULTS

We applied our feature selection algorithm on the relative abundance profiles of 241 microbial species detected in 193 CRC and control samples obtained from the merging of three whole genome shotgun sequencing datasets retrieved through the *curatedMetagenomicsData* package [17] for R: *ThomasAM_2018a, ThomasAM_2018b*, and *ThomasAM_2019_c* (see the Methods section for additional details). Taking in mind the well-known sexual dimorphism in individuals affected by CRC [18] and recent studies that identified some gut-bacteria as possible age-specific biomarkers for the identification of CRC [19–21], we stratified the samples according to biological sex (*male* and *female*) and age category (*adult* and *senior* with age ≤65 and >65 respectively). For each of these datasets, we transformed the species profiles into a binary matrix with 0 and 1 values only, where we consider the presence/absence of microbes independently from their level of relative abundance (RA) (1 if the RA of a particular species in a specific sample is >0, otherwise 0). Feature selection and HD classification model results on binarized and relative abundance datasets are shown in Table 1A.

**Table 1:**
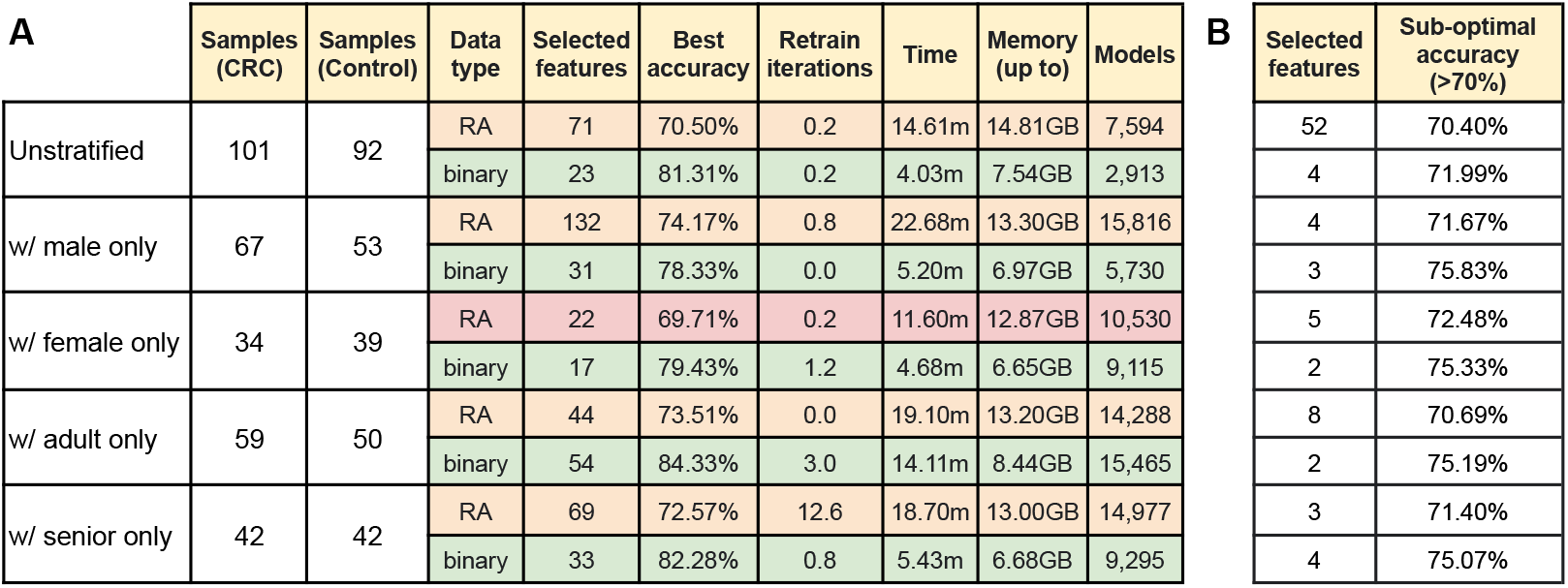
Panel **A**: Feature selection and classification results on (i) the whole dataset resulting from the merging of the *MetaPhlAn3* profile tables of *ThomasAM_2018a, ThomasAM_2018b*, and *ThomasAM_2019_c* (here called *Unstratified*), and the same merged table stratified by biological sex and age category, by considering (ii) male subjects (w/ male only), (ii) female subjects (w/ female only), adult subjects (age ≤65, here called *w/ adult only*), and senior subjects (age >65, here called *w/ senior only*). Every dataset presents the same number of features (241 species) and it is processed twice with relative abundance (RA) and binary presence (binary) microbial profiles, by reporting the number of selected features, the best accuracy, the number of retraining iterations (average on the number of folds) for the classification model with the best accuracy, the computational time used to build and evaluate all the HD classification models up to the end of the feature selection algorithm (in minutes – m), and the amount of allocated memory, in addition to the total number of produced classification models; Panel **B**: Number of features selected by *chopin2* from sub-optimal HD classification models whose accuracy rate is lower than the best one but always >70% with the minimum number of selected features.

| A |  |  |  |  |  |  |  |  |  | B |  |  |
| --- | --- | --- | --- | --- | --- | --- | --- | --- | --- | --- | --- | --- |
|  | Samples (CRC) | Samples (Control) | Data type | Selected features | Best accuracy | Retrain iterations | Time | Memory (up to) | Models |  | Selected features | Sub-optimal accuracy (>70%) |
| Unstratified | 101 | 92 | RA | 71 | 70.50% | 0.2 | 14.61m | 14.81GB | 7,594 |  | 52 | 70.40% |
|  |  |  | binary | 23 | 81.31% | 0.2 | 4.03m | 7.54GB | 2,913 |  | 4 | 71.99% |
| w/ male only | 67 | 53 | RA | 132 | 74.17% | 0.8 | 22.68m | 13.30GB | 15,816 |  | 4 | 71.67% |
|  |  |  | binary | 31 | 78.33% | 0.0 | 5.20m | 6.97GB | 5,730 |  | 3 | 75.83% |
| w/ female only | 34 | 39 | RA | 22 | 69.71% | 0.2 | 11.60m | 12.87GB | 10,530 |  | 5 | 72.48% |
|  |  |  | binary | 17 | 79.43% | 1.2 | 4.68m | 6.65GB | 9,115 |  | 2 | 75.33% |
| w/ adult only | 59 | 50 | RA | 44 | 73.51% | 0.0 | 19.10m | 13.20GB | 14,288 |  | 8 | 70.69% |
|  |  |  | binary | 54 | 84.33% | 3.0 | 14.11m | 8.44GB | 15,465 |  | 2 | 75.19% |
| w/ senior only | 42 | 42 | RA | 69 | 72.57% | 12.6 | 18.70m | 13.00GB | 14,977 |  | 3 | 71.40% |
|  |  |  | binary | 33 | 82.28% | 0.8 | 5.43m | 6.68GB | 9,295 |  | 4 | 75.07% |

It is worth noting that 10,000 was selected for HD vector dimensionality and 1,000 as the number of HD levels for the datasets with RA profiles, while only 2 HD levels were specified for the binary representations of microbial profiles. The maximum number of retraining iterations was set to 10, with retraining terminated when the error rate for an iteration does not vary from the previous round. Here, the accuracy threshold has been set to 60%, while the accuracy uncertainty percentage has been set to 1%. Finally, every classification model was subjected to 5-fold cross validation. The computational time is reported in Table 1A and refers to the total amount of time required for building and evaluating all the HD models generated during the backward variable elimination process. Computation and evaluation of the HD models was performed on a high-performance computing machine comprising 4 Intel Xeon Platinum 8276L CPUs (128 cores/224 threads) @ 2.20GHz and 6TB of RAM using 80 cores at most. Running time and memory consumption profiles have been computed with *memory-profiler* v0.61.0 with an average of 13.43GB of allocated RAM in case of RA models and 7.25GB in case of binary models.

In terms of accuracy, the HD models built on top of the binary datasets (data type *binary*) always reached over 78% of accuracy, which is also always higher than the accuracy of the same models built over the relative abundance profiles (data type *RA*; 74.17% accuracy in the best case, 69.71% accuracy in the worst case).

In Table 1A, we also reported the number of selected features related to the best classification model generated through the iterative steps of our algorithm (see the *Availability* section for the complete list of selected microbial species). It is worth noting that, in order to reach these results, the software generated and evaluated over 32 thousand classification models in total (see column *Models* in Table 1A).

### Sub-optimal classification models significantly reduced the number of relevant features with a minimal loss of accuracy

Here, we also considered sub-optimal results (classification models with lower accuracy rate, always greater than 70%, with the minimum number of selected features) for the binary models, since we noticed that the algorithm extracted a very small number of features able to distinguish the CRC class of samples from the control class of samples with an accuracy slightly lower than the best one. Sub-optimal results are summarized in Table 1B. Since the results about the RA datasets are always worse in terms of accuracy and number of selected features, here we focused on the classification and feature selection results of models built on the binarized datasets only.

The best sub-optimal HD model built on top of the binarized *Unstratified* dataset was able to discriminate the CRC and Control samples with 71.99% of accuracy by considering only 4 features (species). Here we report all of them with their prevalence percentage (*P*) in both CRC and control samples (*P*_*CRC*_ and *P*_*Control*_ respectively): *Eubacterium ventriosum* (*P*_*CRC*_=54.46%; *P*_*Control*_=38.05%), *Firmicutes bacterium CAG 145* (*P*_*CRC*_=53.47%; *P*_*Control*_=29.35%), *Gemella morbillorum* (*P*_*CRC*_=39.6%; *P*_*Control*_=3.26%), and *Parvimonas micra* (*P*_*CRC*_=46.53%; *P*_*Control*_=9.78%).

Despite the feature selection algorithm extracting only 4 species from the *Unstratified* dataset while building a high accuracy classification model, results on the same dataset stratified by biological sex and age category are characterized by an even smaller set of features able to discriminate samples with roughly the same level of accuracy (always >70%).

The algorithm selected 3 features only for the dataset with samples from male individuals with a classification model that reached 75.83% accuracy: *Anaerotruncus colihominis* (*P*_*CRC*_=58.21%; *P*_*Control*_=32.07%), *Bifidobacterium pseudocatenulatum* (*P*_*CRC*_=23.88%; *P*_*Control*_=60.38%), and *Parvimonas micra* (*P*_*CRC*_=44.78%; *P*_*Control*_=7.55%). The number of species selected for the dataset with samples from female individuals is 2, while the achieved accuracy is 75.33%: *Blautia sp CAG 257* (*P*_*CRC*_=14.71%; *P*_*Control*_=35.9%) and *Gemella morbillorum* (*P*_*CRC*_=50%; *P*_*Control*_=2.56%).

The two datasets stratified on the age category are also characterized by a similar situation. Our method selected 2 species for the dataset with samples from adult individuals with a 75.19% accurate classification model: *Oscillibacter sp CAG 241* (*P*_*CRC*_=57.63%; *P*_*Control*_=38%) and *Parvimonas micra* (*P*_*CRC*_=59.32%; *P*_*Control*_=6%). Similarly, the model built on the dataset with samples from senior individuals produced an accuracy of 75.07% with 4 species: *Actinomyces sp HMSC035G02* (*P*_*CRC*_=35.71%; *P*_*Control*_=26.19%), *Bifidobacterium pseudocatenulatum* (*P*_*CRC*_=16.67%; *P*_*Control*_=47.62%), *Streptococcus pasteurianus* (*P*_*CRC*_=11.9%; *P*_*Control*_=0%), and *Streptococcus salivarius* (*P*_*CRC*_=50%; *P*_*Control*_=88.1%).

### Differential abundance analysis of species selected with sub-optimal models confirms known microbial biomarkers

These results show a prevalence of the Firmicutes phyla among all the selected species (9 out of 11). Here we computed the log2 fold change (log2FC) of all the species highlighted by our algorithm as the log2 of the mean relative abundance of the case samples on the mean of the relative abundance of the control samples (mean_CRC_/mean_Control_). Only 4 out of a total of 11 species have an absolute value of log2FC ≥ 1: *Gemella morbillorum* (log2FC=3.6 in *unstratified*, log2FC=4.28 in *female*), *Parvimonas micra* (log2FC=2.25 in *unstratified*, log2FC=3.3 in *adult*), *Bifidobacterium pseudocatenulatum* (log2FC=-1.33 in *male*, log2FC=-1.51 in *senior*), and *Blautia sp CAG 257* (log2FC=-1.28 in *female*).

For these 4 species with absolute value of log2FC ≥ 1 selected from the dataset with unstratified samples, only 2 species belonging to the Firmicutes phylum exhibited statistically significant differential relative abundance according to the Wilcoxon rank-sum test (whose p-values have been corrected for multiple hypothesis testing with the Benjamini-Hochberg procedure): *Gemella morbillorum* (p-value=1.32e-5; FDR=1.59e-3), and *Parvimonas micra* (p-value=1.05e-5; FDR=1.59e-3). These statistically significant species have been selected according to the thresholds of 0.05 on their p-values (p-value ≤ 0.05) and 0.2 on the FDR (FDR ≤ 0.2).

We repeated the same statistical analysis on the other four datasets whose samples have been stratified according to biological sex and age, obtaining the same effect as correcting the Wilcoxon test for these covariates. In the case of the dataset with samples from male individuals only, the 2 selected species that have an absolute value of log2FC ≥ 1 also have statistical significance of differential relative abundance according to the Wilcoxon test and the same threshold of 0.05 and 0.2 on the p-value and the FDR respectively: *Bifidobacterium pseudocatenulatum* (p-value=6.15e-4; FDR=7.42e-2) and *Parvimonas micra* (p-value=4.77e-4; FDR=7.42e-2). For the dataset with samples from female individuals, only one species resulted in statistical significance among the two with an absolute value of log2FC ≥ 1: *Gemella morbillorum* (p-value=5.05e-4; FDR=1.21e-1).

Following the same thresholds, the statistical test highlighted one significant species with log2FC ≥ 1 in the case of the dataset with samples from adult individuals: *Parvimonas micra* (p-value=1.72e-6; FDR=4.16e-4). Finally, there are no statistically significant species for the dataset with samples from senior individuals according to the same threshold used for the other datasets.

According to the most recent updates of the Disbiome database [22], at the time of writing, 2 out of the 3 statistically significant and differentially abundant species identified here are linked to studies in which CRC-affected individuals are involved. Their abundance has already been proposed as possible biomarker for the detection of the disease and are well know to be involved in the destabilization of the colonic wall of the gut, reported as possible cause of progression of CRC: *Gemella morbillorum* [23,24] and *Parvimonas micra* [23,25,26].

Although the Bifidobacterium pseudocatenulatum does not show up in the Disbiome database as linked to CRC, it has also been recently reported as a possible biomarker for the disease [27].

### A comparison with classical wrapper-based feature selection methods

In order to validate our method, we performed a comparative analysis with different feature selection strategies based on Random Forest, Decision Tree, Support Vector Machine (SVM), Logistic Regression, and Extreme Gradient Boosting (XGBoost), on the same binarized datasets in 5-folds cross-validation. Table 2 shows the results in terms of features which have been selected in at least 4 out of 5 folds and the accuracy reached by the classical classification models. Features have been selected according to their importance which produced the classification model with the best accuracy.

**Table 2:** Feature selection based on the importance of the features assigned by specific machine learning models (Random Forest, Decision Tree, Support Vector Machine - SVM, Logistic Regression, and Extreme Gradient Boosting - XGBoost) built on the 5 unstratified and stratified datasets, reporting the number of selected features (*F*), the accuracy (ACC), and running time (*t*) in seconds (s) and minutes (m). Models have been 5-fold cross-validated, and the accuracy (*ACC*) is the mean value among the models with the highest accuracy in all 5 folds. We considered a feature as selected if it turns out to be significant in at least 4 out of 5 folds (columns *Features*). The accuracy levels are color coded according to the difference in accuracy with respect to the *chopin2* results in Table 1 (light red if the accuracy is lower than the accuracy computed on the classification model built with *chopin2*, dark red with NA value if no features were significant in at least 4 out of 5 folds, while no color has been applied in cases where the classical approaches performed better than the level of accuracy computed on the *chopin2* models).

|  | Random Forest |  |  | Decision Tree |  |  | SVM |  |  | Logistic Regression |  |  | XGBoost |  |  |
| --- | --- | --- | --- | --- | --- | --- | --- | --- | --- | --- | --- | --- | --- | --- | --- |
|  | F | ACC | t | F | ACC | t | F | ACC | t | F | ACC | t | F | ACC | t |
| Unstratified | 2 | 80.30% | 16m | 1 | 72.56% | 1s | 6 | 77.70% | 33s | 11 | 81.86% | 11s | 18 | 79.79% | 28m |
| w/ male only | 13 | 83.33% | 28m | NA | 65.83% | 1s | 5 | 81.66% | 44s | 2 | 80.00% | 10s | 5 | 76.66% | 3m |
| w/ female only | 1 | 78.09% | 15m | 1 | 76.57% | 1s | 3 | 76.57% | 36s | 2 | 78.00% | 8s | 1 | 80.85% | 2m |
| w/ adult only | 18 | 89.00% | 16m | 1 | 80.77% | 1s | 1 | 80.64% | 20s | 3 | 87.18% | 9s | 2 | 84.37% | 3m |
| w/ senior only | 2 | 77.50% | 10m | 1 | 72.57% | 1s | 4 | 75.00% | 37s | 4 | 73.82% | 9s | 2 | 70.07% | 2m |

Note that *chopin2* performed better in terms of accuracy compared to most of the classification techniques adopted in this comparative analysis (in average, +2.87% for the *Unstratified* dataset, +0.84% for the *male only* dataset, +1.42% for the *female only* dataset, -0.06% for the *adult only* dataset, and +8.49% for the *senior only* dataset). Also note that in the case of Support Vector Machine and Logistic Regression models, we computed the absolute values on the feature coefficients in order to consider the importance of the features both with negative and positive effects.

Note that 4 out of 4 species selected by *chopin2* in the context of the sub-optimal model built on top of the *Unstratified* dataset were found to be also relevant for at least one out of 5 feature selection techniques applied on the same dataset (i.e., *Eubacterium ventriosum, Firmicutes bacterium CAG 145, Gemella morbillorum*, and *Parvimonas micra*). For the stratified datasets, the number of features selected by *chopin2* that are also significant for at least one of the 5 classical feature selection techniques are: 2/3 (i.e., *Bifidobacterium pseudocatenulatum* and *Parvimonas micra*) and 1/2 (i.e., *Gemella morbillorum*) for the dataset with samples from *male only* and *female only* individuals respectively, and 1/2 (i.e., *Parvimonas micra*) and 2/4 (i.e., *Bifidobacterium pseudocatenulatum* and *Streptococcus salivarius*) for the datasets with samples stratified by age category (*adult only* and *senior only* respectively).

It is worth noting that, although the feature selection technique implemented in *chopin2* relies on the backward variable elimination strategy by iteratively removing features at each step as explained in the Methods section, and thus most of its running time is dedicated to the construction and evaluation of multiple classification models, our approach produced more accurate results compared to the 5 classical machine learning approaches adopted in this analysis. In particular, considering all 5 datasets, *chopin2* generated over 105 thousand models in our experimentation in roughly 2 hours, which is the same time required for running the whole comparative analysis.

The comparative analysis has been performed in Python 3.8 with the support of the *scikit-learn* library v0.22.1 without tuning of methods’ parameters. A complete overview of the results, together with the code to fully reproduce them, is available on Zenodo (see the *Availability* section). It is also worth noting that we have not shown here a comparative analysis on the datasets with microbial relative abundance profiles. We have not discussed these results since the classification models built on the binarized datasets always performed better in terms of accuracy and extracted features. However, classification and feature selection results for these models are also available on Zenodo.

## DISCUSSION

Here, we presented a novel feature selection algorithm based on the backward variable elimination strategy. It has been developed on top of a supervised classifier built according to the hyperdimensional computing paradigm, originally proposed for classifying massive datasets with commodity hardware. To the best of our knowledge, this is the first feature selection method working entirely with vector-symbolic architectures. We want to stress the point that, despite the fact that we tested our method on biological data, the software is domain-agnostic and can be easily applied on data from different knowledge domains. Specifically, we tested our algorithm on abundance profiles of microbes in metagenomic samples of stool in a case/control scenario related to CRC. We identified 11 species able to discriminate case and control samples with high accuracy. Only 3 of them, i.e. *Bifidobacterium pseudocatenulatum, Gemella morbillorum*, and *Parvimonas micra*, have been highlighted by the statistical analysis and are well-known in literature to be linked in some way to the pathogenesis of the disease or as a possible cause of development of CRC. However, 3 of the remaining 8 species, *Eubacterium ventriosum, Streptococcus pasteurianus*, and *Streptococcus salivarius*, are also linked to CRC despite their abundance profiles not showing any statistical significance between case and control samples [28–33]. Finally, there is no mention in the CRC literature about the remaining 5 species (*Firmicutes bacterium CAG 145, Anaerotruncus colihominis, Blautia sp CAG 257, Oscillibacter sp CAG 241*, and *Actinomyces sp HMSC035G02*), and they must be further investigated to determine their role in relation to their abundance as possible rapid and non-invasive diagnostic tool for colorectal cancer in sex- and age-specific individuals.

We are planning to apply the same technique to support large-scale metagenomic analyses in relation to other pathologies, not necessarily limited to cancer diseases. We are confident this will highlight new insights and will help define new highly-accurate non-invasive diagnostic tools for the detection of such pathologies. Additionally, recent advancements in the development of novel techniques for identifying and profiling unknown microbes [34] drastically increase the number of microbial species quantified in metagenomic samples to tens of thousands. Classifying this kind of datasets can be challenging for the state-of-the-art classification and feature selection methods, and the technique we proposed in this manuscript can be a valid solution for efficiently dealing with such a massive number of features. Furthermore, we are also working on improving the overall performance of *chopin2* by running the process of building hypervectors on GPUs and on distributed environments powered by Apache Spark in order to unlock new potentialities of the software for scaling on hundreds of thousands of features.

## MATERIALS AND METHODS

We focused our analysis on the relative abundance (RA) of microbial species profiled over 101 stool samples of patients affected by CRC, in addition to 92 samples from stool of healthy individuals [35,36]. The authors of the selected studies aimed at identifying microbial signatures were able to discriminate the CRC class of samples from the control ones with high accuracy in fecal samples from shotgun metagenomics sequencing. Here we attempt to reproduce the same results with a novel HD-oriented approach. It is worth noting that the 193 samples come from three different datasets (i.e. *ThomasAM_2018a, ThomasAM_2018b*, and *ThomasAM_2019_c*) and their metadata, as well as their microbial abundance profiles originally performed with *MetaPhlAn3* [37], have been obtained through *curatedMetagenomicData* [17] package for R. Table 3 shows a summary of the 193 metagenomic samples involved in this analysis. It is worth noting that 113 out of 193 samples come from antibiotics-free individuals, while for the remaining samples this information is lacking. Furthermore, these three datasets also contain 27 samples of patients affected by Adenoma. However, the number of Adenoma patients is too unbalanced compared to the number of CRC and control samples, thus we excluded them from our analysis.

**Table 3:** Demographic summary of all metagenomic samples involved in this study and retrieved from public datasets. They comprise 193 stool samples distributed over two classes (Colorectal Cancer, and Controls) from female and male subjects ranging from 28 up to 84 years old grouped in two sub-classes (adult and senior) according to the threshold of 65 years old.

|  | Number of samples | Sex (F/M) | Age (years) |  |  |  |
| --- | --- | --- | --- | --- | --- | --- |
|  |  |  | mean (stdev) | range (min–max) | N adult (age ≤ 65) | N senior (age > 65) |
| Colorectal Cancer (CRC) | 101 | 34/67 | 62.41 (11.75) | 28–84 | 59 | 42 |
| Control | 92 | 39/53 | 62.85 (10.53) | 40–81 | 50 | 42 |

We finally looked at the distribution of the samples over the age of the 193 individuals involved in the study in order to check whether the number of samples in a class is too unbalanced compared to the number of samples in the other class and vice versa also in relation to biological sex (*male* and *female*). The distribution charts are shown in Figure 1, Panels A, B, and C. Considering the similar trend of the number of samples from *female* and *male* individuals over the control and CRC classes, we opted to consider all the 193 samples for our study.

**Figure 1:**
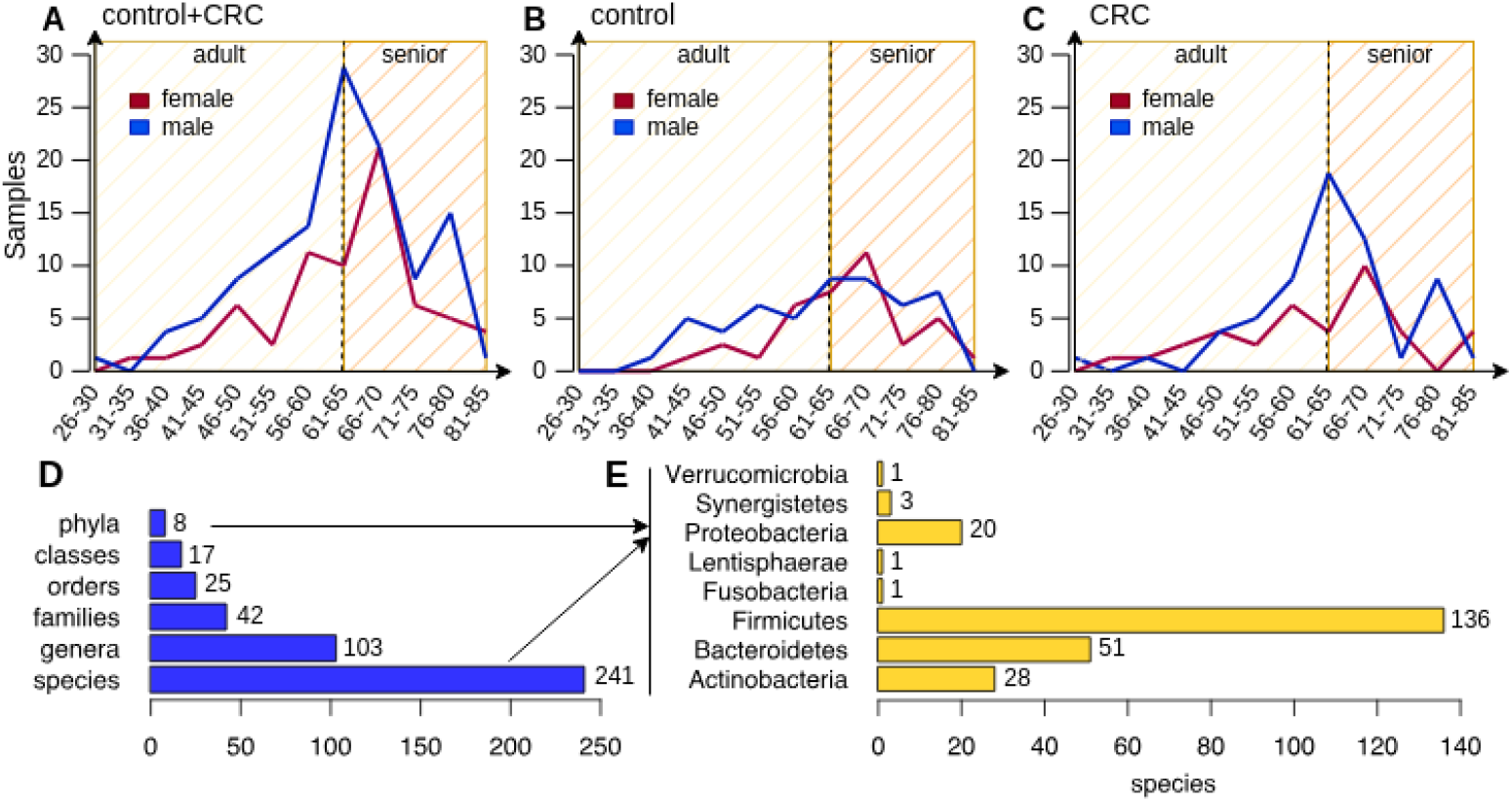
**(Panels A, B, and C)** Distribution of the 193 samples over the control and CRC classes in relation to the age and biological sex of the involved individuals. Panel A shows the overall distribution of all samples, while Panel B and C show the distribution of control and CRC samples respectively. It is worth noting that samples have been grouped according to 5-years age intervals and the vertical dashed line at year 65 in all the three panels represent the time point where an adult turns senior; **(Panels D and E)** Compact view of the microbial composition in the 193 CRC and Control samples. Panel D shows the total number of phyla, classes, orders, families, genera, and species under the Bacteria kingdom, while the barplot in Panel E shows the number of species for each of the 8 phyla detected in the dataset by *MetaPhlAn3*.

### Preprocessing of microbial profiles

Since these samples come from different datasets, we first excluded those microbial species that have not been detected in all the three studies by *MetaPhlAn3*. We can safely exclude these species by simply adding their relative abundances to the *unclassified* profile, which contains the abundance of all the unknown microbial species that *MetaPhlAn3* was not able to detect in a particular sample, resulting in a total of 337 species. We repeated the same process for very low abundant species with a relative abundance lower than 1% (66 species), as well as species detected in less than 5% of the samples (30 species) in order to decrease the possibility of erroneous classifications. This reduced the total number of profiled microbes to 241 bacterial species whose distribution is shown in Figure 1 (241 species, 103 genera, 42 families, 25 orders, 17 classes, and 8 phyla as reported in Panel D, with a majority of species under the Firmicutes phylum (56.4%) as shown in Panel E). It is worth to note that *MetaPhlAn3* reports a relative abundance profile for each of the taxonomic levels, from the kingdom down to the species, for each detected microbe. However, here we focused on the species level only.

### Defining the Vector-Symbolic Architecture and Machine Learning Model

As widely described in previous studies [4,38–40], designing an analysis following the hyperdimensional computing principles means that every atomic element in the input dataset must be first encoded into high-dimensional vectors. Every operation is thus reduced to dealing with a very limited set of arithmetic functions applied on the encoded vectors in the high-dimensional space. The set of vectors and functions is called vector-symbolic architecture (VSA) [4].

We designed the procedure described in Algorithm 1 to build a classification model over a VSA defined on an input numerical dataset:

#### Algorithm 1

Encoding data into high-dimensional vectors

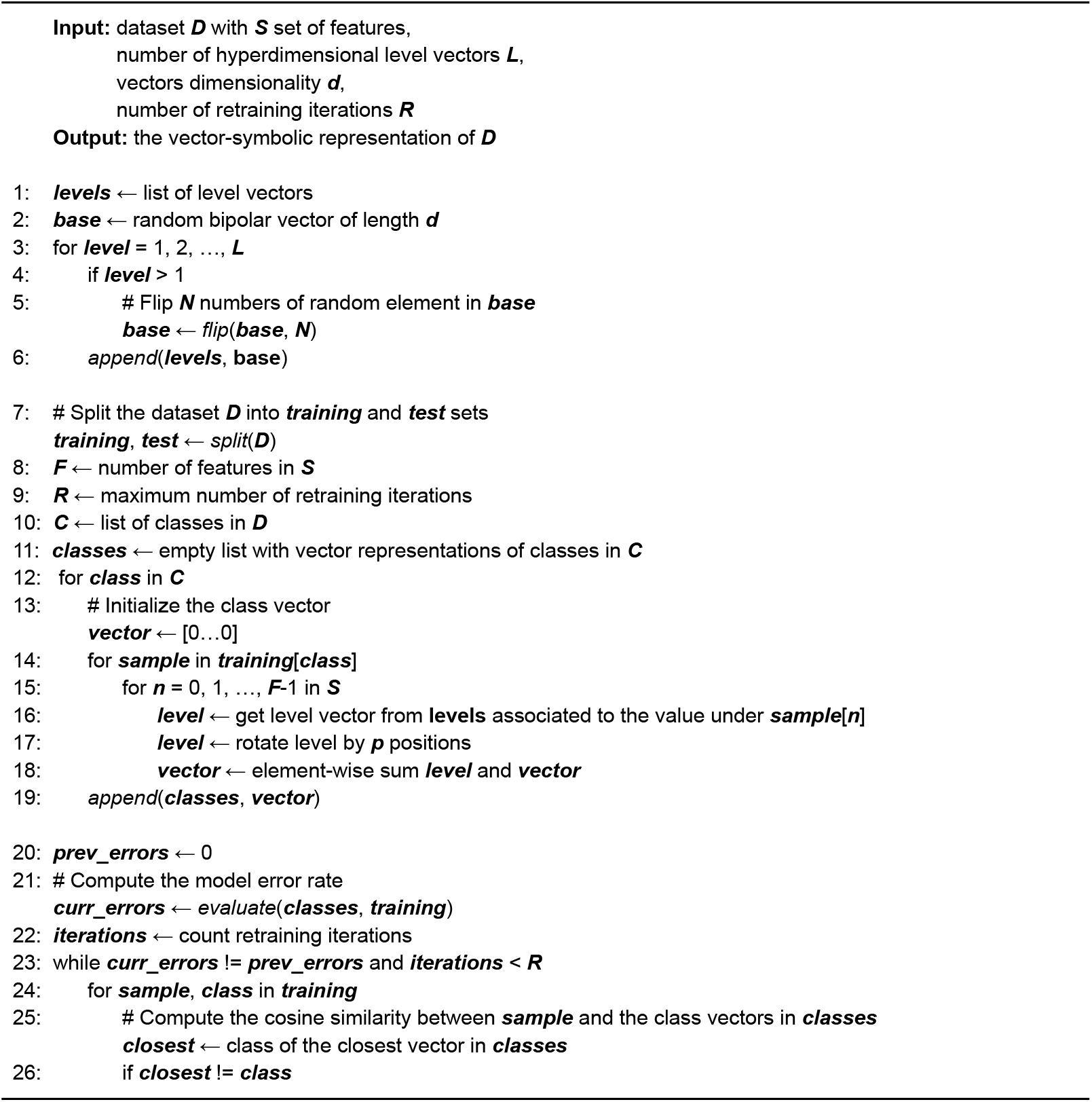
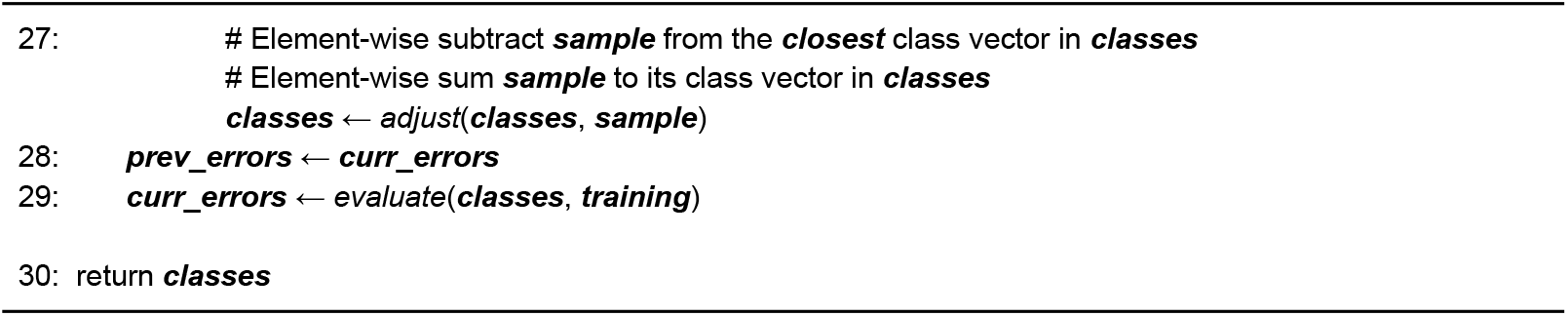

#### Algorithm 1

Pseudocode of the procedure designed to build a vector-symbolic architecture and machine learning model implemented in *chopin2*. It is composed of three main steps: (i) the generation of hyperdimensional level vectors, (ii) the definition of the vector representations of classes whose set represents the model, and (iii) the retraining process (error mitigation) for optimizing the model’s prediction power.

First, it is important to have a look at the distribution of the numerical values in the input dataset. In the case of microbial profiles here analyzed, we know that values range from 0 to 100 because they express relative abundances. This information, as well as the precision of the numerical values, is extremely relevant because every single value must be encoded into a random bipolar vector (containing 1 and -1 values only) in the high-dimensional space (i.e. the length of vectors, usually 10,000). These vectors are usually built starting from a single random vector and iteratively flipping values under a certain amount of random positions (*N* elements are flipped where *N* is equal to the vector dimensionality *d* divided by 2 over the total number of level vectors *L*). This guarantees that adjacent vectors are more similar compared to all the other level vectors by also maintaining their quasi-orthogonality in the high-dimensional space. Vectors must be approximately orthogonal so that they resolve at discernable distance from each other and thus could represent different concepts (e.g., the number 0.002 is different from the number 0.0019, no matter how close these numbers are, and thus they must be encoded into two approximately orthogonal vectors so that they can be easily discriminated just by computing their distance in the high-dimensional space). These vectors are called HD levels and represent the associative memory. The generation of HD levels is the first step towards the encoding of a numerical dataset and it is described in Algorithm 1, lines 1–6. It is worth to note that two different numbers can actually be mapped to the same level vector according to the total number of levels and the global numerical interval in the dataset (e.g., if a dataset contains numbers between 0.0 and 100.0, and the number of level vectors is 100, every number between 0.1 and 1.0 is mapped to the same level vector).

As every classification method usually works, the set of samples in the dataset must be first decomposed into training and test sets (Algorithm 1, line 7). The next step consists in building a vector representation of each class by encoding and combining all the vector representations of the samples in the training set. In order to do that, a sample is first encoded by binding (element-wise sum) all the vector representations of the features under that specific sample. The position of the features is also relevant in defining the encoded vector associated with the samples. In order to also track the feature information, the vector representation of a specific numerical value is rotated (by permuting values) by *n* positions with *n* equal to the position of the feature in the dataset (*n = 0…F-1*, with *F* equal to the number of features in the dataset; the vector representation of the numerical value under the first feature is not permuted, while the vector associated with the numerical value under the second feature is permuted of a single position, and so on up to the last feature in the dataset). The vector representations of the samples belonging to a specific class are finally bounded together for building the vector representation of that specific class. The set of vectors, one for each class, together with the HD levels and this specific way of encoding data, represents the vector-symbolic architecture, i.e., our classification model. The process of encoding data into vectors and combining them to produce the vector representation of classes is described in Algorithm 1, lines 7–19.

The vector representation of a class is thus built by bounding together all the vector representations of samples that belong to that specific class. This operation is extremely easy but summing up vectors also leads to the introduction of noise that affects the prediction power of the whole classification model. We can reduce the inevitable effects of introducing noise with an error mitigation technique known as retraining. This process works by first predicting the class of the training samples that is established by computing the inner-product (cosine similarity) between the vector representation of the training samples and each vector representation of the classes. The closest class is the prediction. In the case of a misclassification, the vector representation of the test sample is element-wise subtracted from the misclassified class vector in order to reduce the noise that led to misclassifying the sample, while it is also element-wise added to the correct class vector to amplify the informative signal of the test sample in the correct class. This process can be performed iteratively many times, usually until the error-rate in a specific iteration converges, i.e., it does not change anymore compared to the previous iteration (Algorithm 1, lines 20–29).

In order to test the model and evaluate its accuracy in correctly predicting the right class, the same process of encoding samples is applied to the test set. The class associated with a sample in the test set is predicted by computing the inner-product of its vector representation and each vector representation of the classes built and adjusted during the previous step. Finally, the accuracy of the classification model is computed as the number of correctly classified samples on the total number of samples in the test set.

### Formalizing the feature selection problem

Here we formally define our feature selection method which belongs to the backward variable elimination class of algorithms. It works by iteratively removing those features that do not have a significant discriminative power for describing observations belonging to a specific class of samples. More specifically, considering a dataset *D* with a set *S* of features, it starts by building a HD classification model considering all the *S* features. The iterative part of the algorithm starts if the accuracy of the model is greater than a specific threshold only.

During the first iteration, it generates all the possible combinations of features by creating *S* sets of *S*-1 features (by excluding a specific feature in all of the *S* sets of features). Then, it builds a HD classification model for each of the *S* sets of features. It keeps track of the features excluded from the specific set of features in case the classification models produced the best accuracy in this specific iteration. Subsequent iterations work in the same way as the first one, except that the initial set of features is now permanently lacking of the previously excluded irrelevant features. It finally stops processing the input dataset when the best accuracy reached in a specific iteration is lower than a predefined threshold (e.g. it may not make sense to keep removing features from a dataset whose classification model has an accuracy <50%) or when there are no other features available in the set of features. Pseudocode for this approach is summarized in Algorithm 2.

#### Algorithm 2

Stepwise Feature Selection as Backward Variable Elimination

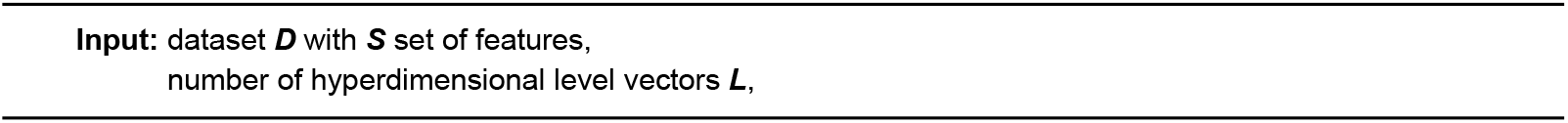
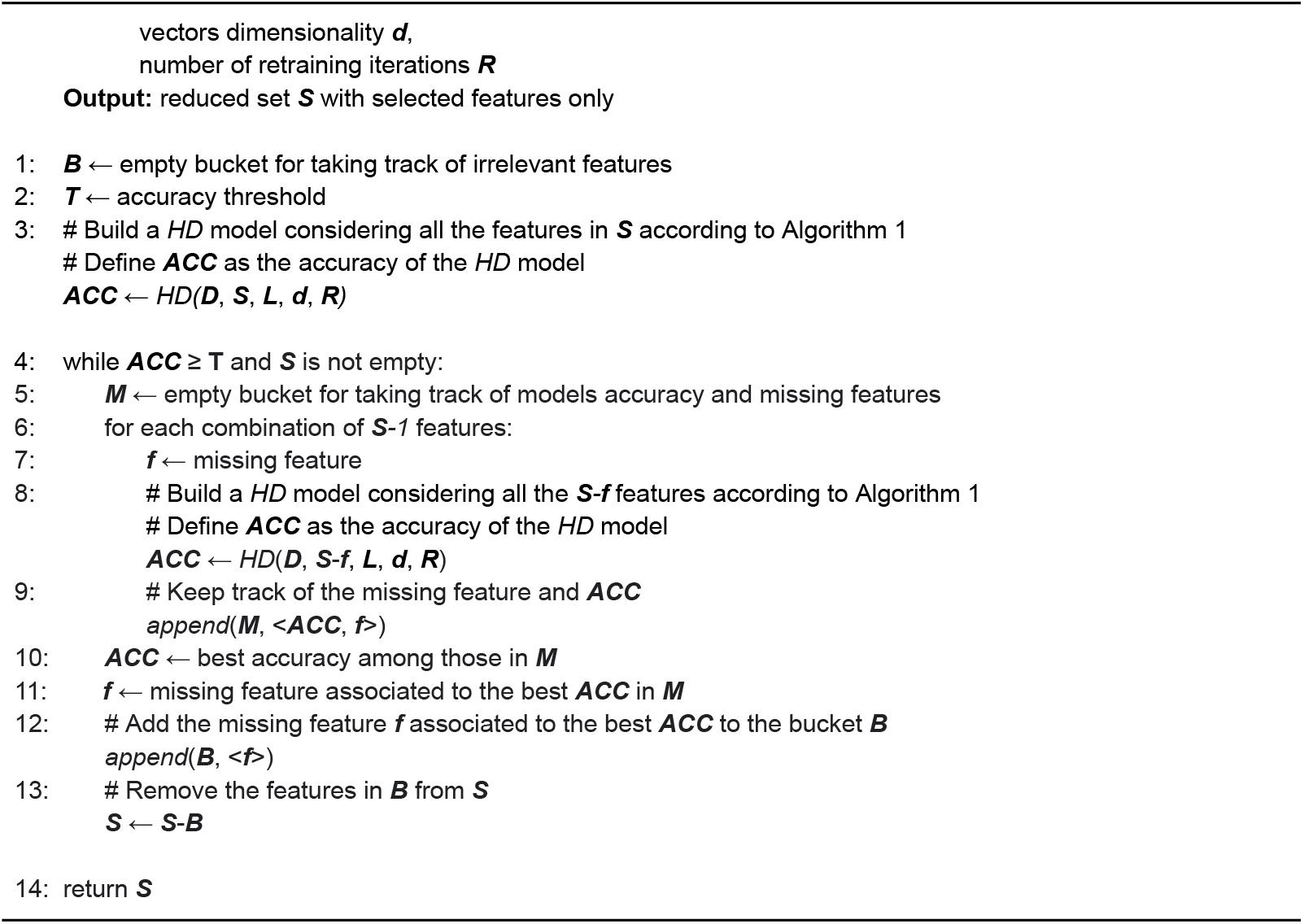

#### Algorithm 2

Pseudocode of the stepwise backward variable elimination algorithm implemented in *chopin2*. Given a dataset *D* in input with *S* features, it iteratively builds a HD classification model by removing irrelevant features that do not contribute to increasing the accuracy of the model itself. It iteratively generates all possible combinations of S-1 features, and it definitively removes the irrelevant feature associated with the model with the best accuracy. It keeps removing features until the best reached accuracy is greater than a specific threshold or when it runs out of features.

To further illustrate this approach, consider the visual representation of the feature selection algorithm in Figure 2. It starts with an example dataset with five features and an empty bucket for taking track of the irrelevant features. It first builds a HD classification model by considering all five features (*S={F1, F2, F3, F4, F5}*) in the input dataset to make sure that there is enough informative content to distinguish the different class of samples with a minimum level of accuracy, here fixed to 70% (step 1). The immediate next step consists in generating all possible combinations of sets of *S*-1 features. Figure 2 (step 2) shows five sets of four features each (i.e. (i) *F2, F3, F4*, and *F5*, (ii) *F1, F3, F4*, and *F5*, (iii) *F1, F2, F4*, and *F5*, (iv) *F1, F2, F3*, and *F5*, and (v) *F1, F2, F3*, and *F4*). For each of them, the algorithm creates a HD classification model that in this particular example produced a 93% as the best accuracy with the second set of features. One of the classification models reached an accuracy of 68%, which is lower than the accuracy threshold of 70%, and thus it must be discarded. Here we also introduced the concept of “*accuracy uncertainty percentage*”, which is a percentage that must be subtracted to the best accuracy in order to define a threshold for establishing whether the accuracy reached by the other models is high enough to be considered as a good result. In this case, the best accuracy is 93% and the “*accuracy uncertainty percentage*” has been fixed to 5%. Thus, the threshold is defined as 88% (93% minus its 5%), and the best results in this iteration are the second and third models with 93% and 91% of accuracy respectively. The algorithm proceeds by applying the same strategy on the third iteration (step 3) by first reshaping the set *S* of features removing *F2* and *F3* which are the features that have been excluded for the generation of the best classification models during the previous iteration. Here, the best accuracy is 90% reached by the last model. There are no other models that must be considered in this iteration since the accuracy reached by the other models is lower than 85% (90% minus its 5%), and in any case lower than the accuracy threshold of 70%. The last iteration (step 4) starts by again reshaping the set of features *S* removing *F5* in addition to *F2* and *F3*. It generates two classification models with only one feature each. Their accuracy is high enough to pass the accuracy threshold, but the best accuracy in this iteration is lower than the best accuracy reached on the previous iteration (step 3). Thus, the algorithm concluded by selecting *F1* and *F4* as the best features able to discriminate the samples in the input dataset with high accuracy, while *F2, F3*, and *F4* do not significantly contribute to increasing the accuracy of the classification model.

**Figure 2:**
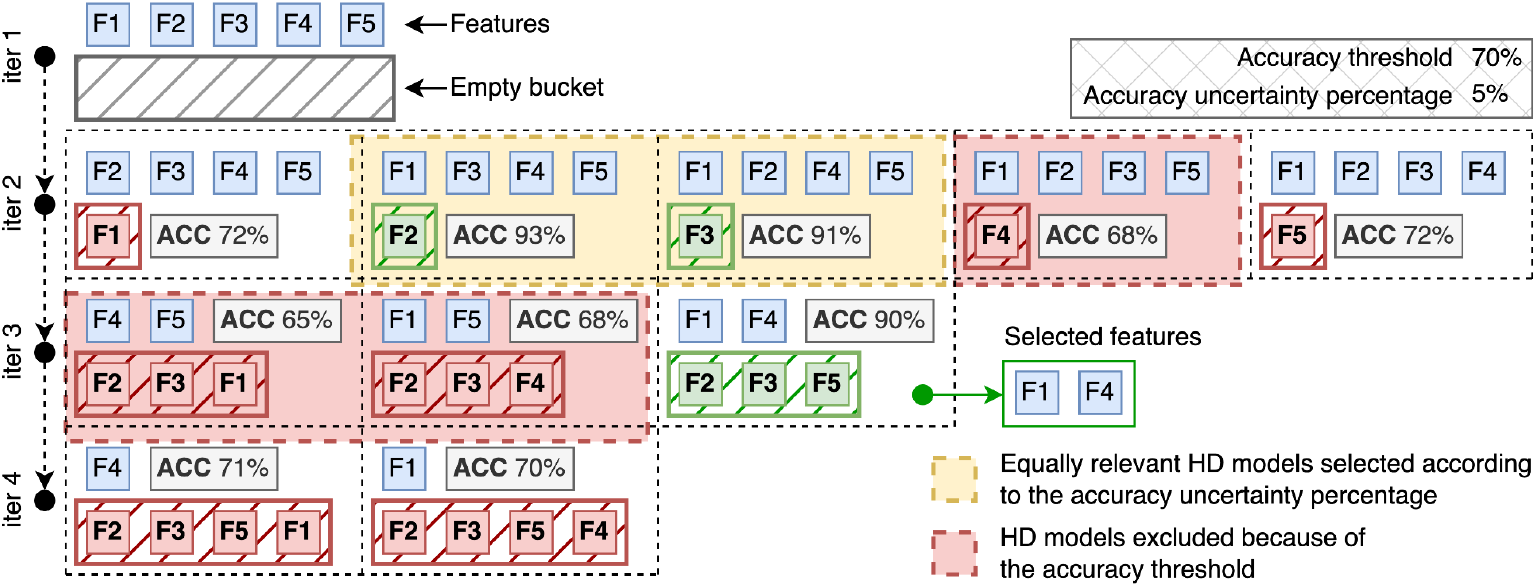
Schematic representation of the backward variable selection algorithm on a dataset with 5 features (F1-5). The first step simply runs the HD classification algorithm on the whole set of features and proceeds if the accuracy of the model is greater than the predefined threshold of 70%. During the second step, five HD models are generated, one for each combination of features by excluding one feature only (i.e., by moving the removed features to the initially empty bucket). Models with accuracy <70% are automatically excluded (red areas), while the excluded features (F2 and F3) of the models with 93% and 91% of accuracy (because 93% is the best reached accuracy and 91% is high enough according to the uncertainty percentage of 5%) are considered to proceed with the third iteration. The algorithm proceeds by generating three models with all the combinations of remaining features minus one. The accuracy of two models does not satisfy the minimum accuracy threshold of 70%, while one model reaches 90% of accuracy with three removed features (F2, F3, and F5). The last iteration produces two classification models because only two features remained active at this point. Their accuracy is way lower than the best accuracy reached by the model at the third iteration (90%). Thus, the algorithm ends by reporting F1 and F4 as the minimum set of features for which the software is able to produce a classification model with high accuracy.

It is worth noting that the algorithm is extremely parallelizable at the single iteration level, since the generation of the HD classification models are completely unrelated from each other. Also note that the same strategy can be easily applied in a reverse order, following a forward variable selection approach, by adding features one by one instead of removing variables. However, this mode is not currently implemented in *chopin2*.

## AVAILABILITY

*MetaPhlAn3* profiles of the analyzed metagenomic samples and their metadata are publicly available through the R package *curatedMetagenomicData*. The proposed algorithm is implemented as a subroutine of the Python 3.8 software *chopin2*, whose source code is available on GitHub under the GPL-3.0 license at https://github.com/cumbof/chopin2, it has been designed through the *hdlib* [41] Python library also available on GitHub at https://github.com/cumbof/hdlib, and it is distributed through the Python Package Index (*pip install chopin2*) and Conda (*conda install -c conda-forge chopin2*). It is also available on the Galaxy platform [42,43] through the official ToolShed [44] at https://toolshed.g2.bx.psu.edu/view/iuc/chopin2 after being revised and approved by the Intergalactic Utilities Commission (IUC). A tutorial for the Galaxy tool is available on the Galaxy Training Network [45–47] at https://gxy.io/GTN:T00337. Classification models and feature selection results presented with this manuscript are fully reproducible and available on Zenodo at https://doi.org/10.5281/zenodo.11397775.

## ACKNOWLEDGEMENTS

We wish to thank the Italian inter-university consortium CINECA for assigning us supercomputing resources (project HDCOMP, grant number HP10CA0SGK to FC) that we extensively used to preliminarily test our algorithm. We would also like to thank Dr. Jayadev Joshi and Dr. Bryan Raubenolt of the Center for Computational Life Sciences, Lerner Research Institute of the Cleveland Clinic, for their constructive critical feedback.

## AUTHOR CONTRIBUTIONS

Conception and design of the study, FC; Software design and development, FC and ST; Methodology and analysis, FC, ST, and EW; Results interpretation, FC and DB; Research supervision, EW and DB; Manuscript preparation FC; All authors provided critical revision of the manuscript and approved the final version for submission.

